# Sex and seasonal variations in melatonin suppression, and alerting response to light

**DOI:** 10.1101/2024.10.18.619012

**Authors:** Fatemeh Fazlali, Rafael Lazar, Faady Yahya, Oliver Stefani, Manuel Spitschan, Christian Cajochen

**Affiliations:** Psychiatric University Clinics (UPK) Basel, Centre for Chronobiology, Basel, Switzerland; University of Basel, Research Cluster Molecular and Cognitive Neurosciences, Basel, Switzerland; Department of Ophthalmology, University of Basel, Basel, Switzerland; Lucerne University of Applied Sciences and Arts, School of Engineering and Architecture, Horw, Switzerland; Technical University of Munich, TUM School of Medicine and Health, Munich, Germany; Max Planck Institute for Biological Cybernetics, Translational Sensory & Circadian Neuroscience, Tübingen, Germany; Technical University of Munich, TUM Institute for Advanced Study, Garching, Germany

**Keywords:** sex differences, seasonal differences, light sensitivity, prior light history, dim light melatonin onset, non-image forming effects

## Abstract

Light influences human physiology and behaviour by regulating circadian rhythms, melatonin secretion, and alertness. Previous research has reported sex differences in melatonin secretion and circadian rhythms, possibly related to women’s greater sensitivity to bright light. Other studies have suggested reduced photosensitivity and earlier circadian phases in summer than in winter in mid-latitude regions. This study explores the effects of sex, seasonality, and their combination on melatonin suppression and subjective sleepiness in response to moderate light exposure, considering prior light history and menstrual phases in females. We conducted a controlled, within-subject experiment with 48 healthy adults (18–35 years, 50% female) across different seasons. The study design included two 9-h laboratory sessions, with at least 5-day washout in between. Participants were exposed to dim and moderate light through a screen for 2 hours after their habitual bedtime. Female participants exhibited greater melatonin suppression (+4.69 %) but a lower alerting response (−6.00%) to moderate light compared to males. Both sexes demonstrated increased sensitivity to the NIF effects of light in winter, with stronger melatonin suppression (+18.05 %) and increased alertness (+7.60 %) compared to summer. While prior light history did not significantly impact melatonin suppression or alertness, it was associated with an earlier dim light melatonin onset (DLMO). Females during their luteal phase had earlier DLMO than those in their follicular phase. Our findings indicate an interaction between sex, seasonality, and light exposure in modulating melatonin suppression, emphasising the need for personalised light exposure recommendations according to individual biological and environmental contexts.

## Introduction

### The non-image forming effects of light (NIF) on physiology and behaviour

Light plays a crucial role in regulating human physiology and neuroendocrine responses^1,2^ through its influence on the circadian pacemaker located in the suprachiasmatic nuclei (SCN) of the brain^3^. The intrinsically photosensitive retinal ganglion cells (ipRGCs)^4–6^ in the eye relay environmental light information directly to the SCN via the retinohypothalamic pathway, thereby synchronising endogenous circadian rhythms with the external 24-hour cycle^7,8^. In addition to its effect on circadian regulation, light exerts acute effects on various aspects of human physiology and behaviour such as the production of the pineal gland hormone melatonin^9–11^, alertness^12,13^, cognition^14,15^, mood, and sleep, which have been described as NIF effects of light^16–18^. Emerging evidence suggests that these NIF effects depend on individual differences in light sensitivity^19–21^, sex^22^ and season^23^.

### Sex differences in the melatonin suppression and alerting response to light

The evidence for differences in light sensitivity between male and female participants is still inconclusive^24,25^, although several studies have shown significant differences^22,26–28^. Female participants typically exhibit an earlier onset of melatonin secretion and higher melatonin concentrations compared to male participants, along with a shorter intrinsic circadian period^29,30^. Moreover, female participants tend to have earlier bedtimes, and wake times, and a higher reported prevalence of sleep issues such as insomnia and daytime tiredness^31–35^. Interestingly, female participants also demonstrate increased sensitivity to bright light-induced melatonin suppression, suggesting a potential mechanism for their earlier circadian timing^22,30,36,37^. These sex differences in light sensitivity appear to be independent of circulating sex hormone levels, as evidenced by the lack of association between melatonin suppression and levels of estradiol, progesterone, or testosterone^22,38^. The inconsistent results in previous studies may be due to methodological differences, such as using different age ranges and not taking the menstrual phase into account^39^. Menstrual phases modulate circadian processes, influencing hormonal rhythms and sleep organisation^40,41^, which could contribute to the variability in light sensitivity observed in female participants. For the sake of clarity, in this paper, we refer to biological sex, not gender, and all participants were cisgender, meaning their biological sex and gender identity were aligned.

### Seasonal variations in melatonin suppression and alerting response to light

Human physiology^42–45^ and behaviour^46^ are affected by the variation in the length of daylight across seasons^47–49^. The SCN responds to changes in photoperiod to adjust melatonin production and sleep timing^50^. Research showing seasonal changes in melatonin secretion patterns^51–55^, circadian rhythms and sleep parameters^49,56–58^ under different light conditions, further supports this regulatory mechanism to explain how seasonality affects human physiology. Both the onset of evening melatonin secretion (i.e. dim light melatonin onset (DLMO)) and the timing of sleep occur earlier^57,59,60^, with shorter sleep duration^58,61^ in summer compared to winter in mid-latitude regions. Extensive research has explored the dynamics between natural and artificial light exposure and their impacts on physiological and behavioural patterns, including sleep^43^. In general, light sensitivity has been found to be higher in winter than in summer^23,63^.

### Our aim

As part of a larger study, the present exploratory analysis examines the effects of sex and seasonality on NIF effects of light on melatonin suppression (area under the curve, AUC; primary outcome) and subjective sleepiness under dim and moderate light exposure levels emitted from a screen display. Initial melatonin data collected in winter showed a clear between dim and moderate light exposure. However, this distinction diminished as we moved into the summer months, prompting an investigation into seasonal variations in melatonin suppression. Thus, inspired by prior work highlighting the need to consider seasonal variables in circadian and sleep research^23^, we conducted a seasonal analysis of our data collected across a year-long period. Furthermore, including menstrual phases in female participants, we explored sex differences in response to light, as sex differences in melatonin suppression have been reported^22^ under bright light conditions (>400 lx) but not under moderate and dim light. Our exploratory analyses extended these observations to more typical light levels, such as those experienced during late evening screen use. Our research focused on exploring the following areas: 1) The potential increased sensitivity of female individuals to moderate light levels compared to male participants, as reflected by lower melatonin AUC under moderate light. 2) The possibility that participants in winter exhibit greater sensitivity to moderate light than those in summer, indicated by higher melatonin suppression during winter. 3) The observed trend that both sexes show heightened light sensitivity in winter compared to summer, with a more pronounced reduction in melatonin AUC. These analyses were conducted in an exploratory manner, providing valuable insights into the observed patterns and guiding our subsequent review of relevant literature. We aim to offer new insights into how light exposure differentially affects individuals based on sex and season, potentially informing personalised strategies for managing circadian and sleep-related problems.

## Materials and methods

### Participants

Forty-eight healthy young participants aged 18–35 years (mean age = 25.11 ± 4.4 years, 50% female, female mean age = 24.99 ± 4.8 years; male mean age = 25.24 ± 4.11 years) with body mass index (BMI) between 18.5 to 29.9 were recruited and participated in our study between October 2022 and October 2023 (winter mean age = 24.9 ± 5.01 years; summer mean age = 25.23 ± 4.15 years). Approximately, twice as many study volunteers participated in the longer photoperiod season defined here as summer season (April to September; n = 31) than in the shorter photoperiod season defined here as winter season (October to March; n = 17). Participant demographics are further detailed in the supplementary information in ***Table S1***. Exclusion criteria included use of any chronic medication affecting neuroendocrine, sleep, circadian physiology, visual system, drugs or nicotine consumption, recent shift work (<3 months), transmeridian travel (>2 time zones) within less than one month prior to the study, high myopia or hyperopia (<-6 and >6 diopters), photosensitive epilepsy, and any ophthalmological issues. Female participants were required not to be pregnant, breastfeeding, or using hormonal contraceptives. Screening was carried out in three phases: online, in-person, and ophthalmological assessments by a physician and an ophthalmologist. The online screening consisted of several questionnaires and tests, including the General Medical Questionnaire (i.e. including vision and hearing health), the ultra-short version of the Munich ChronoType Questionnaire (µMCTQ)^65^, the Pittsburgh Sleep Quality Index Questionnaire (PSQI)^66^, the Epworth Sleepiness Scale (ESS)^67^, the Beck Depression Inventory (BDI-II)^68^, the Alcohol Use Disorders Identification Test (AUDIT)^69,70^ and the Ishihara colour blindness test^71^ using the online tool REDCap^72,73^. Individuals with extreme chronotypes (µMCTQ score: ≤2 and ≥7), poor sleep quality (PSQI score > 5), high depression (BDI-II >13), and moderate alcohol consumption (AUDIT>7) were excluded from the study. In the in-person screening, visual acuity and colour vision tests were conducted using the Cambridge Colour Test (CCT); single Landolt C and Trivector^74,75^, and the Farnsforth 100-hue test^76^ under standardized illumination, which led to the exclusion of participants with colour vision deficiencies (CCT: Trivector; protan >10, deutan>10, and tritan>15, 100-hue error score > 40) or reduced vision (CCT: Landolt C; visual acuity < 0.5). Female participants were tested for pregnancy (i.e. M-Budget pregnancy test, Migros-Genossenschafts-Bund, Zurich). Participants were also excluded if they were unable to understand or comply with the study procedures. Finally, comprehensive physical and ophthalmological screenings were conducted to confirm participants’ eligibility in the study. All inclusion and exclusion criteria are detailed in the supplementary information ***Table S2***.

### Ambulatory phase

Participants underwent an ambulatory phase one week before the study session. During this time, they were instructed to avoid alcohol, irregular caffeine intake, and bananas and citrus fruits (i.e. both on the day of the study sessions) to minimise potential influences on salivary melatonin levels^77^. A consistent sleep-wake schedule was maintained and monitored by wrist actigraphy (Condor Instruments ActTrust 2) and an online sleep diary hosted on REDCap^72,73^. If participants deviated from their bedtime or wake time by more than 30 minutes on any day within the five days prior to each visit, the session was either postponed or the participant was excluded. Menstrual cycle information was also collected because of its possible influence on melatonin concentrations^78^. To ensure consistent physical activity, participants were asked to refrain from strenuous exercise the day before the sessions ^79^. Before each session, drug and alcohol screening was performed using a urine multidrug test (Drug-Screen-Multi 6; nal von Minden, The Hague, The Netherlands) and an alcohol test (ACE X Alcohol test; ACE Instruments, Germany).

### Study protocol

The current study is part of a larger study whose data will be published elsewhere. Here, the study protocol consisted of two 9-hour laboratory sessions, ranging from 6 hours before to 3 hours after the participants’ habitual bedtime (HBT), in constant environmental conditions with no external time cues (***Fig 1a***). All participants were exposed to both dim and moderate light conditions in a within-subject study design. A washout period of at least five days was implemented between the two sessions to avoid any carry-over effects from the previous light condition. For one participant the washout period was 3 days due to the time constraints of her stay in Basel. Identical procedures were applied to all participants in both sessions, the only difference being the 2 hours of moderate light exposure after habitual bedtime. During the control evening session, participants were exposed to dim light (8 lx) for a duration of 7.5 hours, which was used as the baseline condition. On the moderate light exposure evening, participants first spent 5 hours in dim light, followed by 2 hours of moderate light at 100 lx after their HBT. Throughout this period, participants were instructed to keep their eyes open and fixate on the centre of the display, using a chin rest to maintain a stable viewing position. A 20-minute complete dark adaptation period was performed prior to the light exposure. In addition, two 45-minute pupillometry measurements were taken: one before dark adaptation and one after light exposure. The results of these measurements will be reported in a separate publication. Upon arrival, participants were screened for alcohol and urine multi-drug testing and wore a GoPro head strap with an attached light sensor. Saliva samples (>1 mL) and subjective sleepiness ratings (KSS)^80^ were collected every 30 minutes and hourly throughout the sessions, respectively. During the laboratory sessions, participants refrained from using cell phones or laptops and were kept unaware of external time. They were allowed to read, study magazines, books, or other materials, solve crosswords or Sudokus, listen to podcasts, or engage in activities that did not involve self-luminous displays. They were asked to maintain a similar level of activity across all visits. To reduce potential masking effects, participants were given a consistent meal content and calorie intake, consisting of three sandwiches at 1.5, 5.5, and 8.5 hours after arrival at the laboratory. Participants had continuous access to water, and dark sunglasses were provided during restroom visits to prevent unintended light exposure. As the study ended after midnight, participants were given the opportunity to sleep at the Centre for Chronobiology.

### Light exposure

We employed a specially developed multi-primary display^81^ (27-inch Full-HD IPS HP-Display) featuring five LED primaries (i.e. at 430, 480, 500, 550, and 630 nm – ***Fig 1b)***^82^. In the control dim light condition, approximately 3% of each LED primary was activated, while in the moderate light condition, 50% of LED output was activated. Adjustment of these five LEDs provided ∼9 lx_mEDI_ in the dim light condition and ∼ 149 lx_mEDI_ in the moderate light condition (i.e. the irradiance of stimuli is shown in ***Fig 1c***). Calibration procedures were conducted using the JETI spectroradiometer 1501 (i.e. JETI Technische Instrumente GmbH, Jena, Germany) positioned at eye level (i.e. at a height of 120 cm, facing 20 degrees downwards), 65 cm from the centre of the screen. All stimulus conditions, summarised in terms of luminance, illuminance, CCT, colour chromaticity, and photoreceptor excitation, are presented in the supplementary information ***Table S3***.

**Figure 1.**
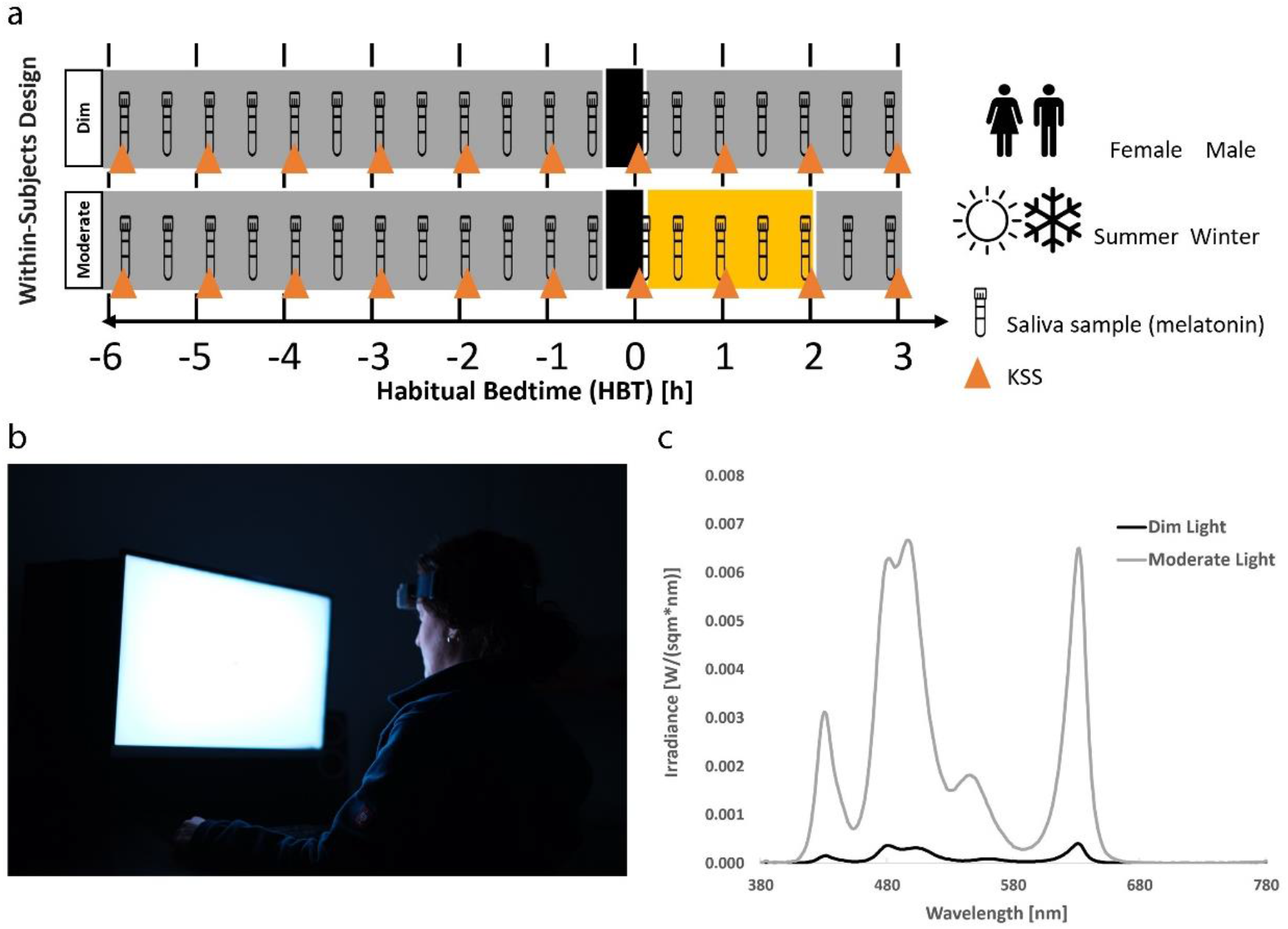
Experimental protocol and lighting conditions (i.e. dim and moderate) of the study. a) For female and male participants in winter and summer. As part of the within-subjects design participants were assigned to both light conditions, 6 hours before to 3 hours after their habitual bedtime (HBT) in randomized order. b) Photo illustrating the screen and experimental conditions. c) Spectral irradiance of dim light (solid black curve; illuminance ∼ 8 lx, mEDI ∼ 9 lx) and moderate light (solid grey curve; illuminance ∼ 100 lx, mEDI ∼ 149 lx). During the control evening, participants were exposed to dim light (≤8 lx) for 7.5 hours, which served as the baseline session. On the moderate-light evening, participants were exposed to dim light for 5.5 hours, and 2 hours of a moderate illumination of (∼100 lx) after HBT. Participants were instructed to fixate on a dot on the centre of the display at all times during the 2 hours of light exposure, using a chin rest. They also underwent a 20-minute period of complete dark adaptation before exposure to the light conditions.

### Salivary melatonin

Salivary samples were collected every 30 minutes during the study session, starting 6 hours before to 3 hours after participants’ HBT. Melatonin levels were measured using a direct double antibody radioimmunoassay (RK-DSM2)^83^. The analytical sensitivity for the detection of melatonin was determined to be at 0.2 pg/mL, with a limit of quantification of 0.9 pg/mL in saliva. All analyses were performed by NovoLytiX GmbH in Witterswil, Switzerland, in accordance with the Instructions of Use of the RK-DSM2 test kit (version 5, 2023-06-08). The area under the curve (AUC) for melatonin during light exposure was computed using the trapezoidal method. The dim light melatonin onset (DLMO) was determined by fitting melatonin profiles with a piecewise linear-parabolic function using the hockey-stick algorithm (version: 2.5)^84^.

### Subjective sleepiness

The one-question 9-point Karolinska Sleepiness Scale (KSS)^85^ was used hourly (i.e. 10 times per session) to assess subjective alerting responses to light. To analyse sleepiness levels, we averaged the KSS ratings for three-time points during the two hours of light exposure for each individual under dim and moderate light conditions.

### Actigraphy and light recordings

To measure individual light exposure and ensure compliance with a regular sleep-wake cycle, participants wore a light dosimeter/actimeter (CAT2; ActTrust 2, Condor Instruments, São Paulo, Brazil) on their non-dominant wrist for at least five days before each study session. Participants were instructed not to cover the device with sleeves. Illuminance data were recorded in 1-min intervals throughout the study period, and pre-processing was conducted using the LightLogR package^86^ in R. We filtered out illuminance data below 1 lx to exclude values from nighttime periods or may have been covered by clothing^87^. For prior light history analysis, we used the duration of time above 100 lx threshold (TAT100 lx) on the day preceding the study session, from wake-up time in the morning to arrival at the laboratory in the evening (∼10 hours). This metric, consistent across both seasons, was the best predictor of melatonin AUC, and a threshold for melatonin suppression and alertness in previous research^88,89^. It should be noted that light exposure at eye level may differ from that measured at the wrist. To investigate the influence of illuminance changes in winter and summer, we illustrated the average 24-h logarithmic illuminance values across days before the study sessions.

### Menstrual phases

We collected menstrual cycle data from female participants during online screening. Female participants were asked to report the start and end dates as well as the length of their menstrual cycles. To estimate the menstrual phase, we used the forward-counting method^90^ based on their self-reported dates. In this approach, we counted the days starting from the onset of menstruation.

### Statistics and reproducibility

All statistical analyses were conducted in R (Version 4.3.0, R Core Team, 2023). We included the melatonin concentration of all participants and calculated the area under the curve (AUC) for melatonin data during two hours of light exposure. The linear mixed model (LMMs) analyses were performed in R using the lme4 packages. The melatonin AUC was included as the main response in the LMM. Light condition, prior light history, and menstrual phases (only females) were included as fixed effects, and repeated measures per participant were modelled as a random intercept in different sexes and seasons. LMMs were followed by an ANOVA (Type III) function. As an effect size measure, partial omega squared (ωp^2^) was calculated using the effectsize package with the following interpretation: small effect: ωp^2^ ≥ 0.01, medium effect:ωp^2^ ≥ 0.06, large effect: ωp^2^ ≥ 0.14. A p-value < 0.05 was considered to indicate statistically significant differences. Linear regression models were used to describe the relationship between DLMO and prior light history in different sexes and seasons and the relationship between DLMO and menstrual phases (follicular and luteal) in female participants. We detected melatonin onsets of all participants’ melatonin profiles during the dim light exposure using the hockey-stick method. We chose a default upper limit or threshold of 2.3 pg/ml for the area of interest and selected ‘Hockeystick Time’. In eleven cases, the threshold had to be adapted as the minimum melatonin values were slightly more than 2.3 pg/ml. Two participants were excluded from the DLMO analyses because we could not detect a melatonin onset in one profile and in another case, the DLMO occurred during light exposure. We adjusted the upper limit of the area of interest to a value that is the first increasing melatonin value below the threshold. The LMMs were also used for the effect of light conditions on the average of subjective sleepiness. We used the same analysis procedure and LMM structure as for melatonin AUC to estimate the average subjective sleepiness (during 2 hours of light exposure). Percentages were calculated using the formula: [(A_Dim_ −A_Moderate_ /A_Dim_)_[female or winter]_ −(A_Dim_ −A_Moderate_ /A_Dim_)_[male or summer]_]×100. This formula was applied to different sex and season groups, where A represents either the melatonin AUC or the average sleepiness. For analyses of the full data set, and subgroup analysis see the supplementary information in ***Table S4-8***.

## Results

We investigated the effects of a screen with a moderate light level on melatonin suppression and subjective sleepiness in the late evening. We also examined the contribution of prior light history as a covariate to these effects. In addition, we analysed the effect of prior light history on DLMO. To explore the effect of menstrual phases on melatonin AUC and DLMO, we included menstrual phases as an additional covariate for female participants.

### Melatonin suppression and alerting response to light

We evaluated the evening temporal profile of melatonin concentration and melatonin AUC during the 2-h exposure under two different light conditions (***Fig. 2a and b***). There was no significant effect of moderate light on melatonin suppression. When assessing the effect of prior light history on melatonin AUC, neither had a significant effect.

**Figure 2.**
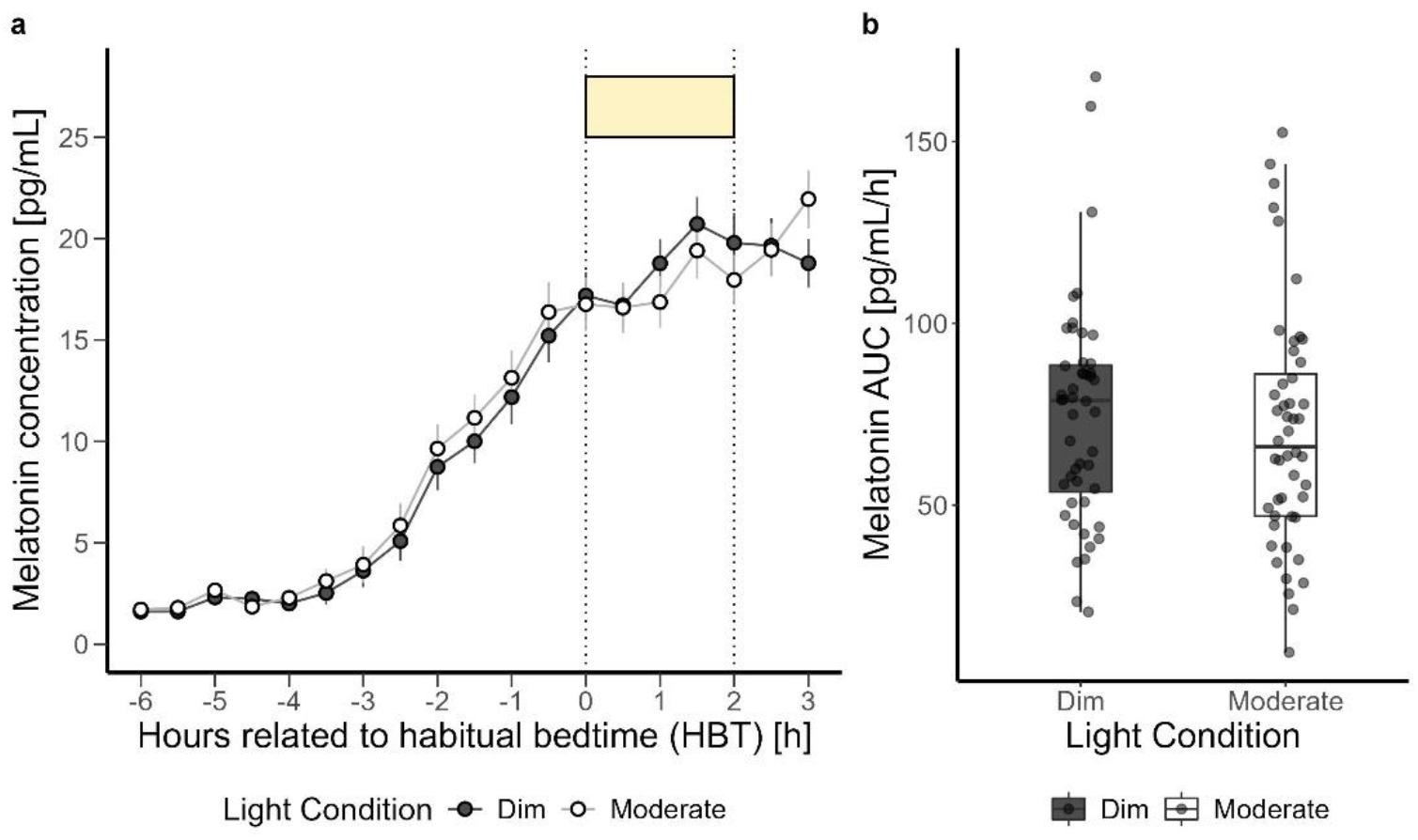
a) Time course of salivary melatonin concentrations (mean ± SEM per 30-min time bins) in pg/mL during dim light (black dashed) and moderate light (grey solid) conditions plotted against the hours relative to habitual bedtime (HBT) in h. b) Area under the curve (AUC) of melatonin in pg/mL/h during the 2-h exposure to dim and moderate screen light. The box plots display the median (central horizontal line), interquartile range (edges of the box, from first quartile; 25^th^ percentile to third quartile; 75^th^ percentile), and the range of minimum and maximum values within 1.5 times the interquartile range from the first and third quartiles (whiskers). Grey dots represent the individual values of participants.

However, the effect of prior light history on DLMO was significant. Results indicated that an increase in prior light history was correlated with an earlier DLMO (F_1,44_ = 8.76, p = 0.004, large effect) (***Fig. 9a***).

Average subjective sleepiness levels showed a typical increase during the late evening, which intensified after the participants’ habitual bedtime (***Fig. 3a***). We observed a significant alerting effect of the moderate light compared to the dim light (F1,46 = 5.41, p = 0.02, medium effect) (***Fig. 3b***), whereas the prior light history did not significantly affect subjective sleepiness during 2-h light exposure. Supplementary ***Table S4*** provides a summary of the analysis for all outcomes. [Fig 3]

**Figure 3.**
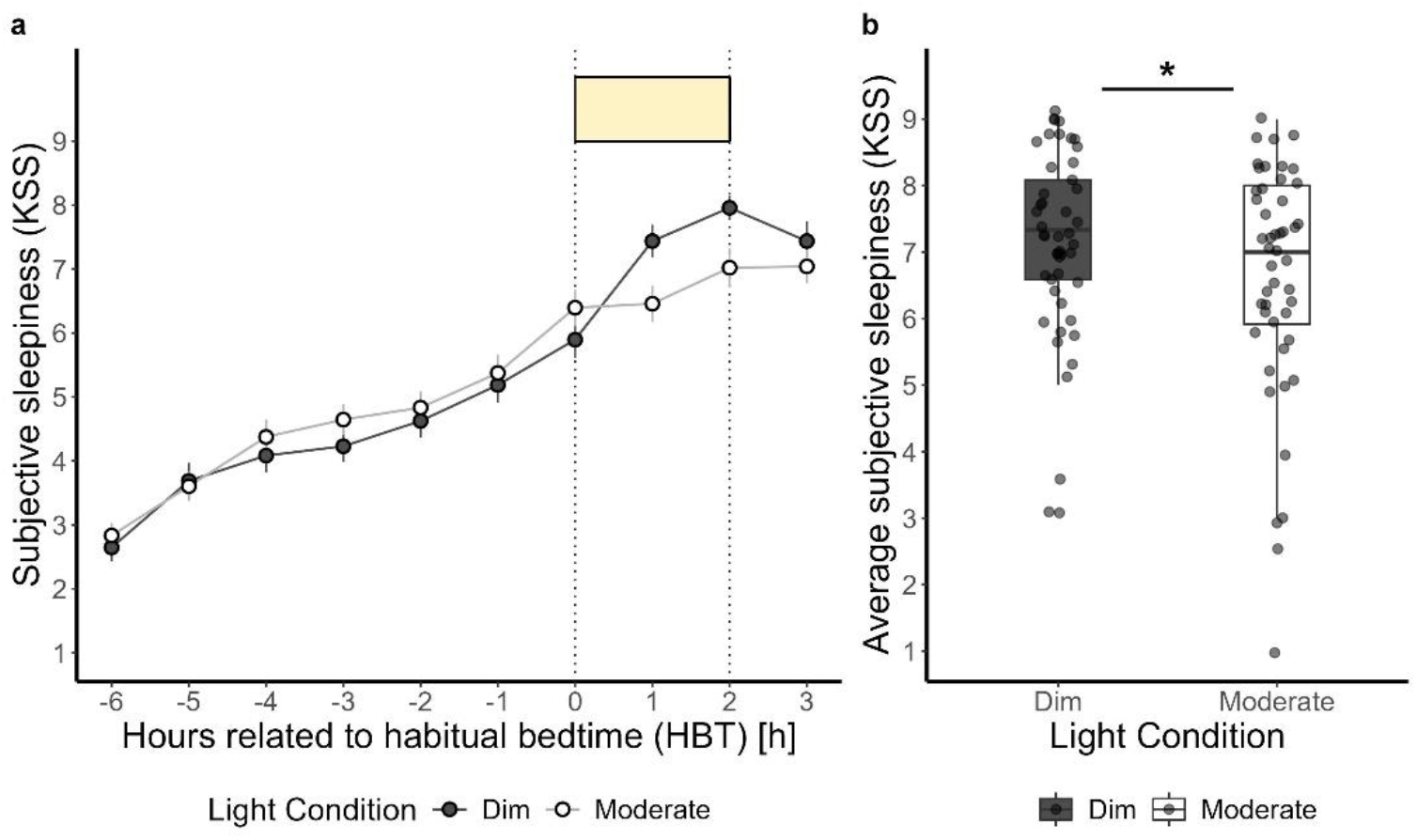
a) Time course of subjective sleepiness ratings (KSS) (mean ± SEM per 60-min time bins) during dim light (black dashed) and moderate light (grey solid) conditions plotted against the hours relative to habitual bedtime in h. b) Subjective sleepiness average over three-time points during the 2-h light exposure for dim and moderate light conditions. Values shown are means of three-time assessments. The box plots display the median (central horizontal line), interquartile range (edges of the box, from first quartile; 25^th^ percentile to third quartile; 75^th^ percentile), and the range of minimum and maximum values within 1.5 times the interquartile range from the first and third quartiles (whiskers). Grey dots represent the individual values of participants. The black asterisk indicates the statistically significant main effect (p < 0.05) of light condition on the 2-h average of subjective sleepiness, calculated in a linear mixed model.

### Sex differences in the melatonin suppression and alerting response to light

Incorporating sex as a fixed effect to the linear mixed model analysis, the effect of light condition on melatonin AUC was not significant (approaching significance p = *0*.*06*). In addition, the impact of prior light history, sex, and the interaction of light condition and sex on melatonin suppression was not significant (Supplementary ***Table S5***). However, female participants showed higher suppression of melatonin under moderate light compared to male participants (+4.69%) (***Fig. 4a and b***). By including menstrual phases in the sex subgroup analysis of female participants, we did not find a significant effect of light condition, as well as prior light history and menstrual phase on melatonin suppression (Supplementary ***Table S6***). [Fig 4]

**Figure 4.**
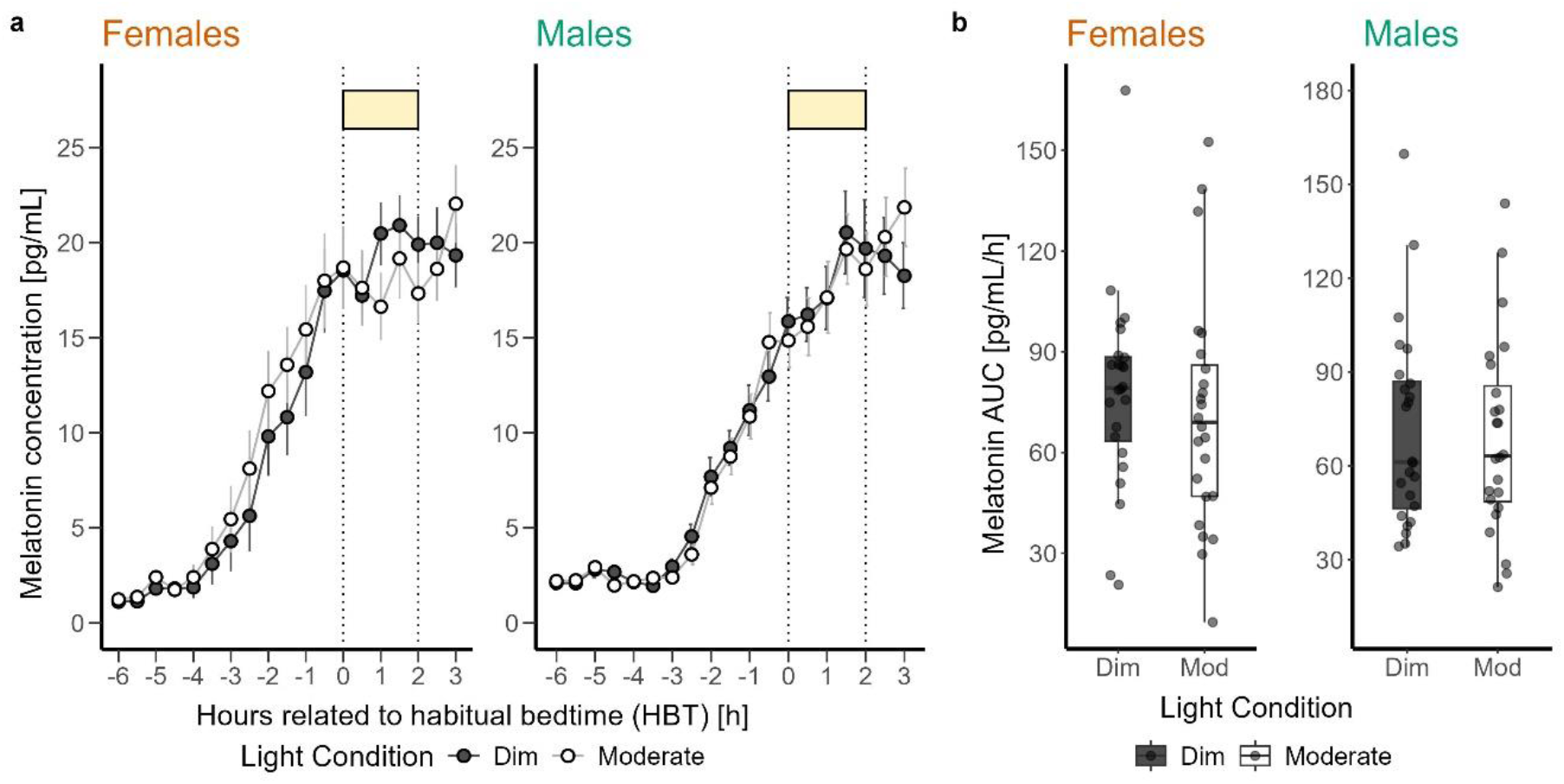
a) Time course of salivary melatonin concentrations (mean ± SEM per 30-min time bin) in pg/mL during dim light (black dashed) and moderate light (grey solid) conditions plotted against the hours relative to habitual bedtime in h in female and male participants. b) Area under the curve (AUC) of melatonin in pg/mL/h during the 2-h light exposure. The box plots display the median (central horizontal line), interquartile range (edges of the box, from first quartile; 25^th^ percentile to third quartile; 75^th^ percentile), and the range of minimum and maximum values within 1.5 times the interquartile range from the first and third quartiles (whiskers). Grey dots represent the individual values of participants.

However, the prior light history significantly influenced the DLMO in both sexes (F_1,43_ = 11.48, p = 0.001, large effect). Increased prior light exposure was associated with earlier DLMO (***Fig. 9b***). There was no significant effect of sex on DLMO. Although female participants on average showed half an hour (∼28 min) earlier DLMO compared to male participants. Moreover, female participants in their luteal phase had earlier DLMO than females in their follicular phase (F_1,21_ = 8.15, p = 0.009, large effect) (Supplementary ***Table S6***).

Average subjective sleepiness profile showed higher levels of sleepiness in female participants compared to male participants in both light conditions before the light exposure (− 7.90 %) (***Fig. 5a***). When sex was included as a fixed effect in the analysis, the effects of light condition on subjective sleepiness was significant with a more alerting response to moderate than dim light (F_1,45_ = 5.42, p = 0.02, medium effect) (***Fig. 5b***). However, the impact of prior light history, as well as, sex and the interaction of light condition and sex were not significant (Supplementary ***Table S5***).

**Figure 5.**
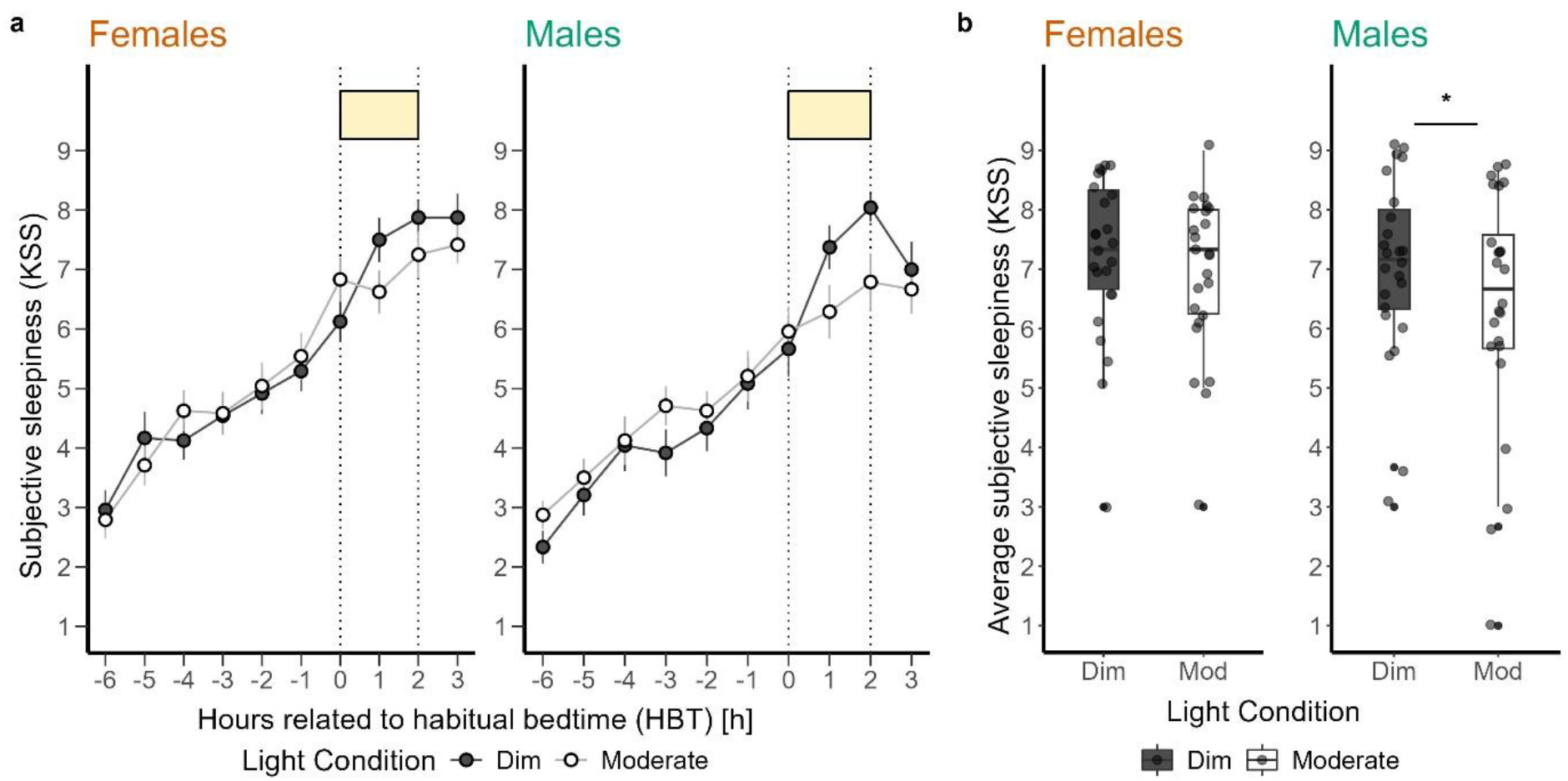
a) Time course of subjective sleepiness ratings (KSS) (mean ± SEM, 60-min time bins) during dim light (black dashed) and moderate light (grey solid) conditions plotted against the hours relative to habitual bedtime in h in female and male participants. b) Subjective sleepiness averaged over three-time points during the 2-h light exposure for dim and moderate light conditions in female and male participants. The box plots display the median (central horizontal line), interquartile range (edges of the box, from first quartile; 25^th^ percentile to third quartile; 75^th^ percentile), and the range of minimum and maximum values within 1.5 times the interquartile range from the first and third quartiles (whiskers). Grey dots represent the individual values of participants. The black asterisk indicates the significant main effect (p < 0.05) of light condition on the mean subjective sleepiness level during light exposure, calculated in a linear mixed model.

### Seasonal variations in the melatonin suppression and alerting response to light

The average logarithmic illuminance levels were significantly higher in summer than in winter (F_1,46_ = 32.94, p < 0.001, large effect) (***Fig 6a and b***). The notable drop in illuminance levels during winter, started at 18:00 and was followed by a stable light exposure until 21:00. There was a significant interaction between illuminance and season (F_1,6_ = 17.58 p < 0.01, large effect). The average logarithmic illuminance levels for different sexes are shown in ***Figure S1*** in the supplementary information.

**Figure 6.**
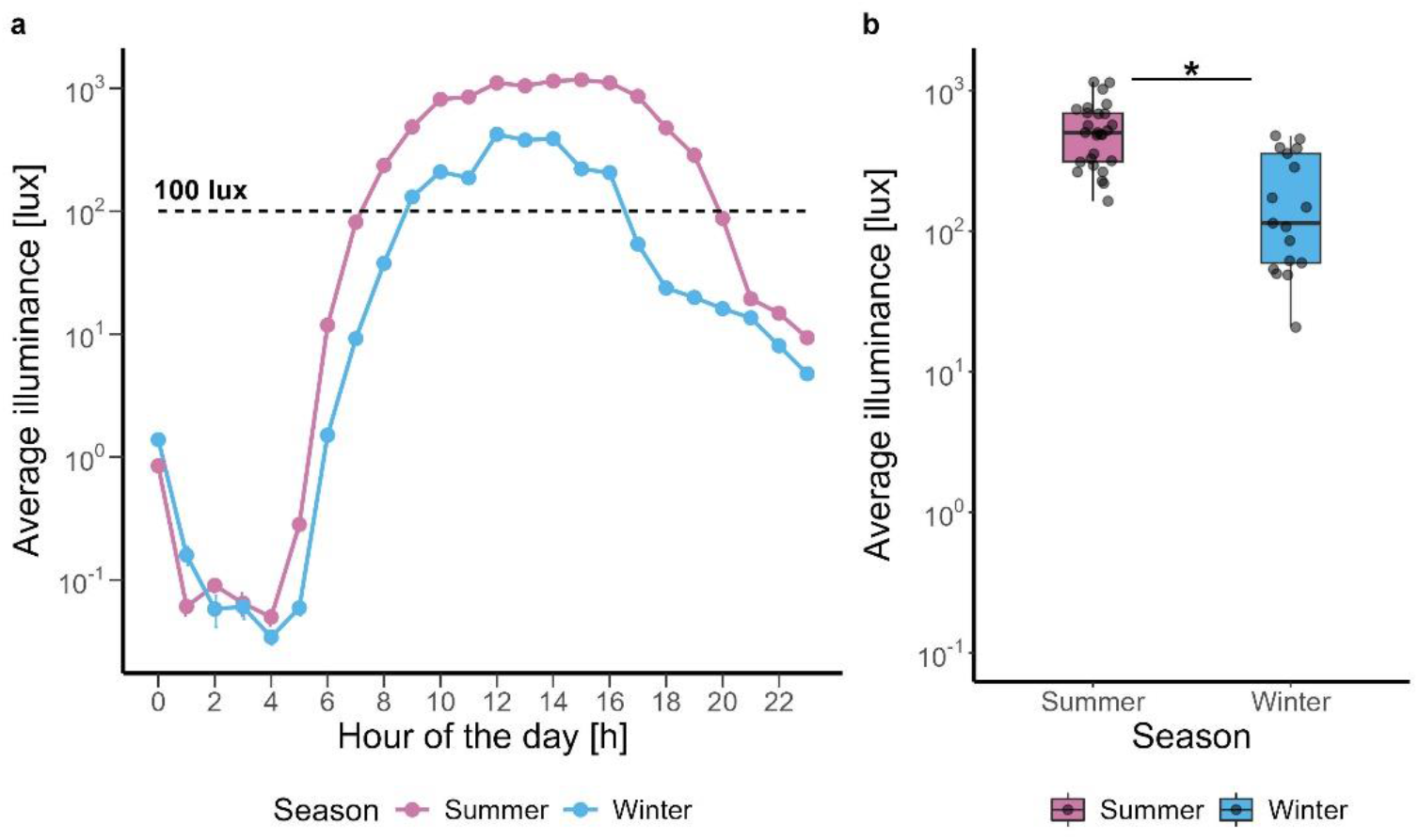
a) Average logarithmic (base 10) illuminance in lx plotted against hour of the day in summer (pink) and winter (blue). b) Average logarithmic illuminance in lx in summer (pink) and winter (blue). The box plots display the median (central horizontal line), interquartile range (edges of the box, from first quartile; 25^th^ percentile to third quartile; 75^th^ percentile), the range of minimum and maximum values within 1.5 times the interquartile range from the first and third quartiles (whiskers), and the outliers (solid dots outside of the whiskers). Grey dots represent the individual values of participants. The black asterisk indicates the significant main effect (p < 0.05) of season on the prior light history, calculated in a linear mixed model.

The average melatonin concentration profile for both seasons revealed that individuals in winter exhibited greater sensitivity to moderate light during the 2-h light exposure compared to summer participants (+18.05 %) (***Fig. 7a and b***). When season was included as a fixed effect in the linear mixed model analysis, the AUC showed significantly greater melatonin suppression in moderate compared to dim light (F_1,45_ = 7.99, p = 0.006, medium effect). Additionally, the interaction of light condition and season was significant (F_1,45_ = 9.34, p = 0.003, large effect). However, the impact of other variables comprising prior light exposure and season did not reach statistical significance (Supplementary ***Table S7***).

**Figure 7.**
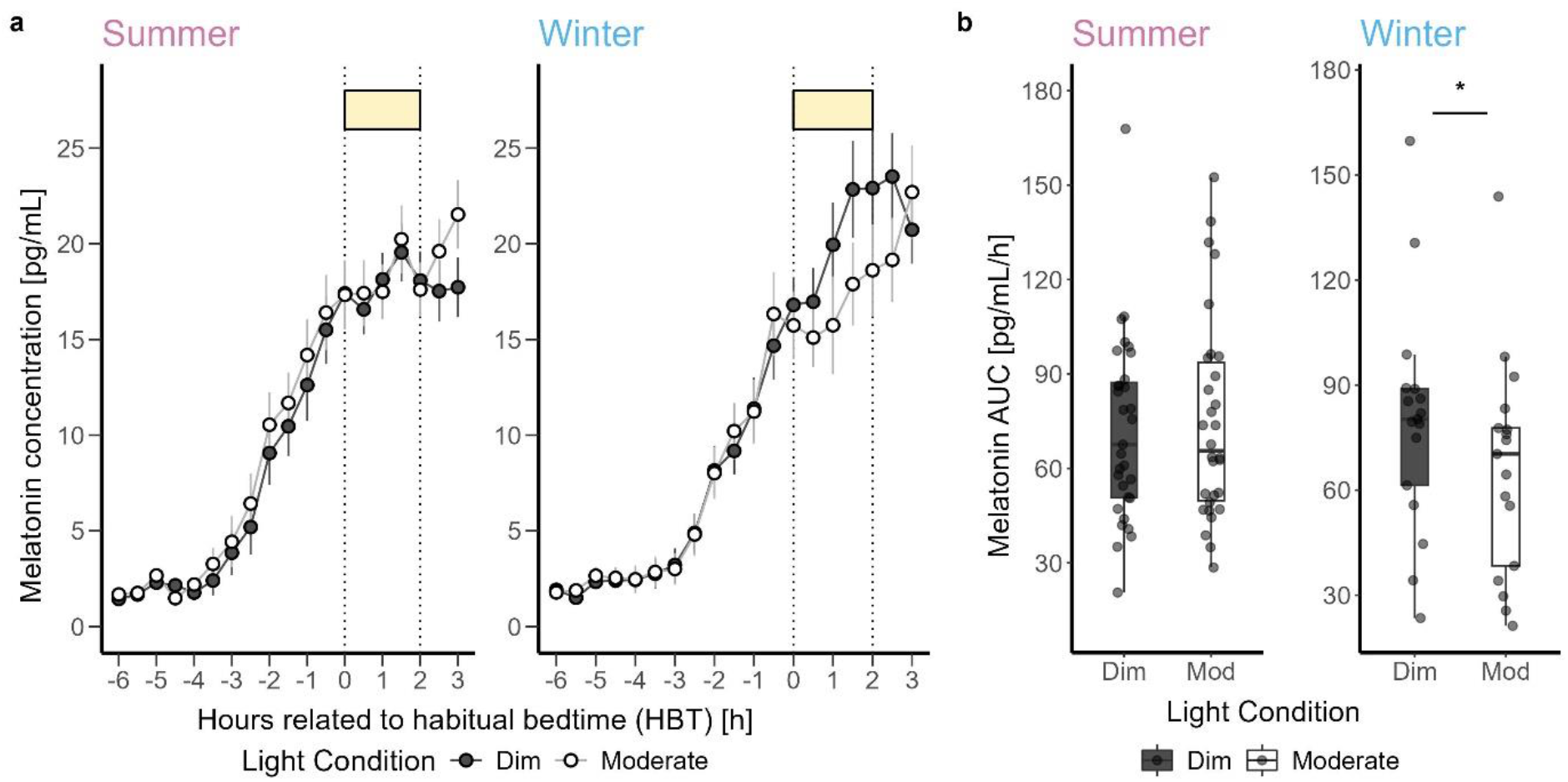
a) Time course of salivary melatonin concentrations (mean ± SEM, per 30-min time bins) in pg/mL during dim light (black dashed) and moderate light (grey solid) conditions plotted against the hours relative to habitual bedtime in h in summer and winter. b) Area under the curve (AUC) of melatonin in pg/mL/h during the 2-h light exposure in summer and winter. The box plots display the median (central horizontal line), interquartile range (edges of the box, from first quartile; 25^th^ percentile to third quartile; 75^th^ percentile), and the range of minimum and maximum values within 1.5 times the interquartile range from the first and third quartiles (whiskers). Grey dots represent the individual values of participants. The black asterisk indicates the significant main effect (p < 0.05) of the fixed factor light condition, calculated in a linear mixed model.

Further analysis showed that prior light history significantly modulated DLMO in both seasons. Participants showed that higher prior light history was associated with earlier DLMO (F_1,43_ = 8.75, p = 0.005, large effect). There was no significant effect of season on DLMO (***Fig 9c***).

The average subjective sleepiness profile illustrated a higher sensitivity to moderate light in winter compared to summer season (+7.60 %) (***Fig. 8a***). Incorporating the season variable into the analysis, the effect of light conditions on subjective sleepiness was still significant with more alerting response to moderate than to dim light (F_1,44_ = 7.03, p = 0.01, medium effect) (***Fig 8b***). However, the effects of other variables were not significant (Supplementary ***Table S7***).

**Figure 8.**
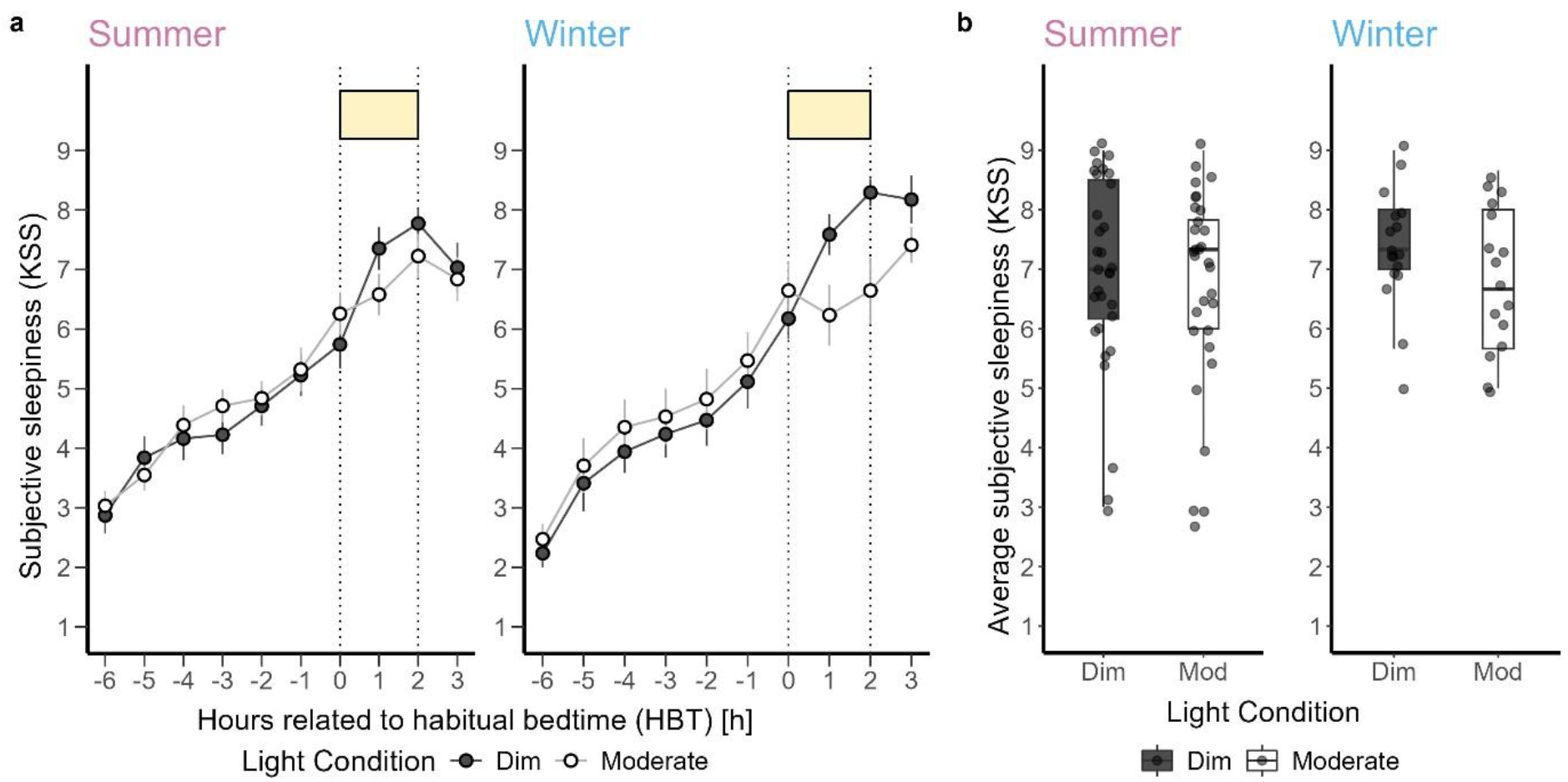
a) Time course of subjective sleepiness ratings (KSS) (mean ± SEM, 60-min time bins) during dim light (black dashed) and moderate light (grey solid) conditions plotted against the hours relative to habitual bedtime in h in summer (left) and winter (right). b) Subjective sleepiness averaged over three-time points during the 2-h light exposure for dim and moderate light conditions in summer (left) and winter (right). The box plots display the median (central horizontal line), interquartile range (edges of the box, from first quartile; 25^th^ percentile to third quartile; 75^th^ percentile), and the range of minimum and maximum values within 1.5 times the interquartile range from the first and third quartiles (whiskers). Grey dots represent the individual values of participants.

**Figure 9.**
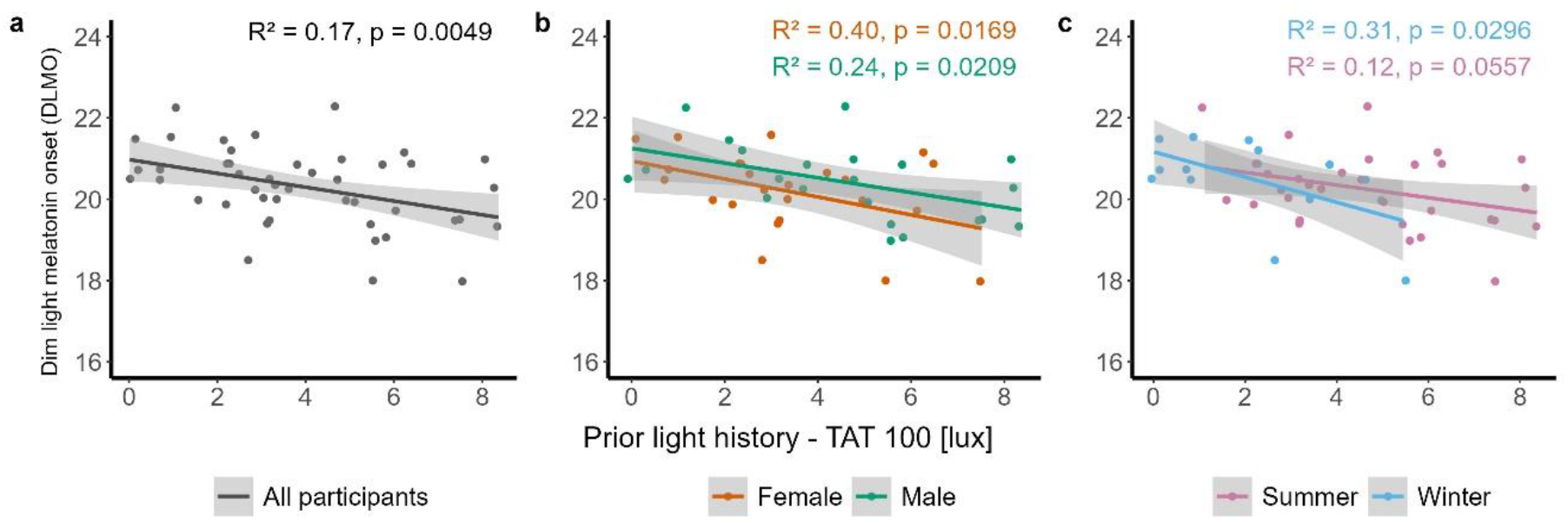
Dim light melatonin onset (DLMO) in hour of the day versus prior light history-duration of time above threshold (TAT) 100 lx in h for a) all participants b) female participants (orange) and male participants (green) c) in winter (blue) and in summer (pink). Dots represent the individual values of participants. The grey band represents the 95% confidence interval limits in the regression model.

### Interactions between sex, season, and light on melatonin suppression and alertness

By including both sex and season as fixed effects in the analysis, the melatonin AUC revealed a significantly higher melatonin suppression under moderate than dim light (F_1,43_ = 8.67, p = 0.005, large effect). The interaction of light conditions and season (F_1,43_ = 10.06, p = 0.002, large effect), and the interaction of sex and season (sex*season; F_1,44_ = 4.39, p = 0.04, medium effect) were significant (Supplementary ***Table S8***). Furthermore, the effect of prior light history on DLMO was significant (DLMO; F_1,41_ = 11.21, p = 0.001, large effect). However, there was no significant effect of sex and season on DLMO. In female participants, the luteal phase was correlated to earlier DLMO (F_1,12_ = 5.87, p = 0.03, large effect) only in summer.

Enhanced alertness under moderate light conditions was observed for both female and male participants during both seasons. The effect of light condition on subjective sleepiness was significant with more alerting response to moderate light (F_1,43_ = 7.28, p = 0.009, medium effect). However, the impact of other variables (i.e. sex and season) was not significant (Supplementary ***Table S8***).

## Discussion

Here we report evidence that in a well-controlled laboratory study probing photoreceptor-specific cone contribution to melatonin suppression, both sex and season modulate, melatonin suppression and the alerting response to moderate light levels. We also found that prior light history influences the circadian phase as indexed by the DLMO. Female participants were more sensitive to light-induced melatonin suppression than male participants but showed a less alerting response to moderate light during 2 hours of light exposure starting after habitual bedtime. Moreover, melatonin suppression by moderate light was stronger during winter than summer for both sexes. The degree of melatonin suppression observed in the laboratory was unrelated to the amount of light female and male participants had been exposed to before the start of the study (i.e. prior light history) in winter and summer. However, the effect of prior light history significantly impacted DLMO for both sex and season groups, with earlier DLMOs in participants exposed to more light prior to the laboratory study. We also found the first evidence that female participants in the luteal phase of their menstrual cycle had earlier DLMO than those in the follicular phase of their cycle.

### Influence of sex on light-induced melatonin suppression and alerting effects at night

Female participants exhibited greater sensitivity to moderate light exposure (149 lx mEDI), showing increased melatonin suppression compared to male participants. This finding is consistent with earlier research by Monteleone et al.^36^ and Vidafar et al.^22^, who reported sex differences in melatonin suppression under different light conditions. In contrast to our findings, they only showed greater sensitivity to bright light (i.e., < 400 lx) but not to moderate light in female than male participants. Our findings also diverge from a previous study by Nathan et al.^25^ who observed no significant sex differences in melatonin suppression at 200 lx. These discrepancies may be attributed to variations in sample size, light intensity, light spectral composition, the timing of the exposure, or the methodologies used to assess light sensitivity. Monteleone et al. conducted their study with 12 participants exposed to 2000 lx of light from 2 am to 4 am, independent of the participants’ habitual bedtime, allowing them to sleep from 11 pm to 2 am and 4 am to 7 am. Vidafar et al. exposed 56 participants to a range of light illuminances (10, 30, 50, 100, 200, 400, and 2000 lx) for 5 hours, 4 hours before to 1 hour after their habitual bedtime. Nathan et al. conducted two different studies: one with 43 participants exposed to 200 lx, and another with 11 participants exposed to 500 lx, both involving light exposure for 1 hour between 12 am and 1 am, regardless of their habitual bedtime. The more pronounced sensitivity to light observed in female participants could be linked to an earlier melatonin secretion onset^29,37^, which was not significant in our data (i.e. an average of 28 minutes), or to a shorter intrinsic circadian period^91^, which we were unable to measure in our study. These biological factors may predispose female participants to a more pronounced melatonin suppression under artificial light conditions in the evening and at night. In contrast to other studies, female participants in our study demonstrated notable differences in melatonin suppression already at light levels, akin to those typical during evening screen use. However, this was not translated into a significantly stronger alerting response to moderate light in female than male participants, aligning with Cowan et al. s’^28^ observation of sex differences in the visual cortex response to light. They demonstrated that male participants exhibited a significantly greater BOLD signal increase in response to blue light compared to female participants, suggesting differential neural processing between sexes under specific light conditions. Another study by Chellappa et al^27^ also found that female participants were less able than male participants to discriminate subjective brightness perception between blue-enriched and non-blue-enriched light exposures. They found that males showed higher brightness perception and faster reaction times in sustained attention tasks under blue-enriched light compared to non-blue-enriched light, whereas female participants did not exhibit significant differences. In our study, male participants reported significantly higher alertness under moderate light levels than female participants, despite weaker melatonin suppression. Our analysis did not reveal a relationship between melatonin suppression and menstrual phase in female participants, echoing findings by Nathan et al^38^. and Vidafar et al.^22^. This suggests that the observed sex differences in light sensitivity may be influenced by other factors, such as genetic variations or differences in photoreceptor cells.

### Influence of season on light-induced melatonin suppression and alerting effects at night

Our exploratory analyses revealed seasonal variations in light sensitivity, with an increase in melatonin suppression in winter compared to summer. This finding is consistent with an earlier study by Wehr^51^, who clearly showed that the circadian profile of human melatonin rhythms changes during simulated long-(i.e. summer) and short (i.e. winter) photoperiod conditions in the laboratory. Complementary findings^23,55,63^ suggest that light sensitivity tends to be greater in winter across different latitudes, likely due to prolonged darkness and reduced daylight exposure, which supports the observations made by Thorne et al.^47^ and Shochat et al.^92^. Thorne et al. analysed daylight exposure from 09:00 to 21:00, while Shochat et al. examined the full 24-hour light profile. Both studies found seasonal variations, with light exposure peaking in the morning and declining more in the afternoon, particularly in winter. Our results aligned with these findings, showing significantly higher light exposure in summer compared to winter and a notable drop in late afternoon light levels during the winter months. As expected, individual daily light exposure measured across five days before the in-lab part of our study started was brighter and of longer duration in summer than in winter. During winter, illuminance levels started to drop around 18:00 and remained stable thereafter until 21:00, demonstrating the influence of artificial lighting on human photoperiodic experience. Moreover, the average subjective sleepiness was higher during dim light and lower under moderate light levels in winter than in summer, which is most likely consistent with higher light sensitivity in winter seasons. This observation partially aligns with the results of Meyer et al^93^, who reported seasonal variations in task-related brain responses, highlighting the significant impact of seasonality on cognitive functions.

### Prior light history and the DLMO

We examined whether the differences in melatonin suppression and alertness between winter and summer could be attributed to significant variations in prior light exposure experienced during these seasons. Our findings indicate no significant effect of prior light exposure on melatonin suppression and subjective alertness. However, a significant effect was observed on the circadian melatonin phase. Longer prior light exposure before the laboratory sessions was associated with an earlier DLMO in both sexes. In line with this, previous studies^59,60^ reported earlier DLMO and time of sleep during summer compared to winter, with delays in sleep timing noted during the winter months, as discussed by Danilenko et al. and Duster et al.^53,57^. The earlier occurrence of DLMO in summer further suggests that seasonal variations in circadian timing might be driven by increased sensitivity to the phase-advancing effects of bright morning light. This underscores the role of prior light exposure history in influencing circadian phase adjustments, suggesting that both immediate and cumulative light exposure histories are crucial in determining our circadian rhythms. Additionally, our findings indicate that the menstrual cycle’s effects on DLMO are observed only during summer, particularly during the luteal phase, earlier DLMO associated with the luteal phase. This partially aligns with Kauppila et al.’s^52,94^ observation on higher levels of Luteinizing hormone (LH) during the luteal phase in summer but not in winter. They also reported higher melatonin levels during the follicular phase in winter, suggesting that higher melatonin levels in winter may inhibit LH secretion. Conversely, lower melatonin levels in summer may reduce this inhibitory effect. However, it should be noted that Kauppila et al. did not investigate DLMO in their studies. This highlights the complexity between light exposure, seasonal changes, and hormonal activity across different menstrual phases.

### Interactions of sex and season on light-induced melatonin suppression and alerting effects at night

Our study demonstrated interactions among sex, seasonality, and light exposure in influencing melatonin suppression. Moderate light exposure resulted in significantly higher melatonin suppression during winter for both female and male participants, which aligns with the increased light sensitivity during winter season. However, within this interaction, the effect of the season was more pronounced than the effect of biological sex. Regarding subjective alertness, no significant interactions among sex, season, and light conditions were observed. These findings underscore the necessity of considering the combined influences of biological sex and environmental factors, such as seasonal variations, to comprehensively understand the effects of light on the human circadian system.

### Limitations

While our study offers valuable insights into the variations in light sensitivity across sex and seasons, it is important to acknowledge certain limitations. The sample size used, though sufficient for identifying significant effects, may limit the generalizability of our findings to a broader population. To enhance the robustness of these results, future studies should include larger and more diverse cohorts. Moreover, our research, focusing on dim and moderate light exposures typical of evening screen use, underscores the need to investigate a broader range of light intensities and spectra to fully understand the spectrum of human light sensitivity and its implications for circadian and sleep regulation. In addition, the analysis of sex and seasonal effects involved between-subject comparisons, which may introduce variability in the results.

## Strengths

This study has several strengths that contribute to its robustness and reliability. First, the use of a well-controlled laboratory setting allowed precise manipulation and measurement of light exposure, ensuring high internal validity. The within-subject design minimised individual variability, thereby increasing the accuracy of our findings on light sensitivity. In addition, the study’s comprehensive screening process, which included several stages and a range of health assessments, ensured the inclusion of a healthy and homogeneous group of participants. Furthermore, the precise measurement of salivary melatonin using a sensitive radioimmunoassay provided accurate and reliable melatonin profiles.

## Conclusion

This study highlights the importance of personalizing light exposure recommendations to improve sleep quality and circadian rhythm disorders. Our findings suggest that in addition to considering light scenarios, taking into account the individual’s sex and seasonal changes could make light-based treatments more effective for sleep and mood issues. This approach is supported by individual differences in the human circadian system and light sensitivity, as explored in previous studies^19,20^. In summary, our study extends the understanding of non-image forming (NIF) effects of light by illustrating how these responses are modulated by a combination of both sex and season.

## Acknowledgements

We are particularly thankful to our medical doctors, Christian Epple and Corrado Garbazza, for their invaluable assistance with the medical examinations. We also wish to acknowledge Jakob M. Weber and Alejandro Fernandez Estrada from NovoLytiX for their expertise in the melatonin assay and Christine Blume for sharing the equipment and laboratory. Special thanks to Johannes Zauner for providing the LightLogR package, which was very helpful for analyzing light history data. We thank all participants who volunteered for this project. The success of this research was made possible by the dedicated efforts of our interns and study helpers: Sonja Camenzind, Rebecca Frommherz, Maria Vettiger, Fiona Vogel, Lisa Tran, Taifun Süner, Nadja Tschan, Bela Bernasconi, Milica Trailović, Karoline Edrich, Melanie Schmid, and Joshua Reese.

## Data and Code Availability Statement

The de-identified data and code for this study will be made publicly available on GitHub and FighShare under a specified licence upon publication.

## Funding Statement

This study is funded by the European Union’s Horizon 2020 program, part of the European Training Network, under the Marie Skłodowska-Curie grant agreement No. 860613 (LIGHTCAP). Moreover, the completion of this work was supported by the “Dissertationen und Habilitationen” grant of Freiwillige Akademische Gesellschaft, the “Financial Contribution” grant of Josef and Olga Tomcsik-Stiftung, and the “Simone and Jacqueline Bühler-Fonds” grant of GGG Basel.

## Conflict of Interest Statement

F.F., R.L., and F.Y. declare no conflict of interest related to lighting. O.S.is named as an inventor on the following patents: (“Display system having a circadian effect on humans”, US8646939B2; “Projection system and method for projecting image content”, DE102010047207B4; “Adaptive lighting system”, US8994292B2; “Projection device and filter therefor”, WO2006013041A1; “Method for the selective adjustment of a desired brightness and/or colour of a specific spatial area, and data processing device”, WO2016092112A1). O.S. is a member of the Daylight Academy, Good Light Group and Swiss Lighting Society. O.S. has had the following commercial interests in the last two years (2022–2024) related to lighting: Investigator-initiated research grants from SBB, Skyguide, and Porsche. M.S. is listed as an inventor on a patent application (“Determining metameric settings for a non-linear light source”, WO2020161499A1). C.C.’s commercial interests between 2022 and 2024 related to lighting include honoraria, travel, accommodation and/or meals for invited keynote lectures, conference presentations or teaching from Toshiba Materials, Velux, Firalux, Lighting Europe, Electrosuisse, Novartis, Roche, Elite, Servier, and WIR Bank.

## Ethics Approval Statement

The study was conducted at the Centre for Chronobiology in Basel, Switzerland, from October 2022 to October 2023. Ethical approval for the study was obtained from the Ethics Commission of Northwest and Central Switzerland (EKNZ) under the approval number 2022-00401, classified as an “other clinical trial”. All experimental procedures adhered to the principles outlined in the Declaration of Helsinki. Participants were provided with a comprehensive information sheet for better understanding and willingly signed a written consent form before engaging in the study. All participants were compensated for their time and participation in this study.

## Patient Consent Statement

Informed consent was obtained from all subjects involved in the study. Written informed consent has been obtained from the participants to publish this paper.

## Clinical Trial Registration

This study was conducted according to the protocol ID 2022-00401. The trial was registered at ClinicalTrials.gov with the identifier NCT05423002.

## Author Contributions

C.C. acquired funding for the study. F.F., R.L., O.S., M.S. and C.C. conceived and designed the experiment. F.F. implemented the test setup and configured the light conditions, supported by O.S., M.S., and C.C. F.F. performed the research and acquired the data. F.F., R.L., and C.C. analyzed and interpreted the data. F.F. drafted the manuscript. F.F., R.L., F.Y., O.S., M.S., and C.C. provided a critical review of the manuscript. All authors have approved the final version of this manuscript.

## Supplementary Information

The online version contains supplementary material available at …

