## Supplementary material for "Sex and seasonal variations in melatonin suppression, and alerting response to light": Table S1-S8, Figure S1

Table S1) participants' characteristics in different sex and season. Data are expressed as mean ± SD. BMI: Body Mass Index; BDI-II: Beck Depression Inventory; µMCTQ: Ultra-Short Version of the Munich ChronoType Questionnaire; PSQI: Pittsburgh Sleep Questionnaire; ESS: Epworth Sleepiness Scale; Visual Acuity with single Landolt C; Farnsworth Munsell 100 Hue test; Cambridge Colour Test Trivector; Protan, Deutan, and Tritan

|  | | | Sex | | Season | |
| --- | --- | --- | --- | --- | --- | --- |
| Descriptives | | | **Female** | **Male** | **Winter** | **Summer** |
| Age | | Mean | 24.99 | 25.24 | 24.9 | 25.23 |
|  |  | sd | 4.8 | 4.11 | 5.01 | 4.15 |
| BMI | | Mean | 22.67 | 22.22 | 22.59 | 22.37 |
|  |  | sd | 2.81 | 1.79 | 2.25 | 2.42 |
| BDI_II | | Mean | 3.67 | 2.42 | 2.94 | 3.1 |
|  |  | sd | 4.06 | 2.95 | 3.29 | 3.76 |
| µMCTQ | | Mean | 3.92 | 4.29 | 4.12 | 4.1 |
|  |  | sd | 0.89 | 0.73 | 0.8 | 0.85 |
| PSQI | | Mean | 3.58 | 2.83 | 3.12 | 3.26 |
|  |  | sd | 1.41 | 1.4 | 1.58 | 1.39 |
| ESS | | Mean | 6.54 | 4.92 | 5.24 | 6 |
|  |  | sd | 3.05 | 2.67 | 2.28 | 3.27 |
| Visual Acuity | | Mean | 1.88 | 1.94 | 1.93 | 1.9 |
|  |  | sd | 0.45 | 0.36 | 0.42 | 0.4 |
| Farnsworth | | Mean | 19 | 17.33 | 18.82 | 17.81 |
|  |  | sd | 11.33 | 9.26 | 9.46 | 10.82 |
| CCT | **Protan** | Mean | 5 | 4.29 | 4.55 | 4.69 |
|  |  | sd | 1.8 | 1.33 | 1.46 | 1.71 |
|  | **Deutan** | Mean | 4.86 | 4.47 | 4.28 | 4.88 |
|  |  | sd | 1.53 | 1.64 | 1.53 | 1.59 |
|  | **Tritan** | Mean | 6.38 | 6.41 | 6.12 | 6.54 |
|  |  | sd | 2.62 | 2.51 | 2.4 | 2.64 |

Table S2) inclusion and exclusion criteria for initial screening survey, in-person screening, physician screening, and every visit screening

| Aspect | Assessment modality | Exclusion criterion and cut-off | Timing of screen |
| --- | --- | --- | --- |
| Age | Self-report | <18 years >35 years | Initial screening survey |
| BMI | Self-reported height and weight | <18.5  >29.9 |  |
| Pregnancy (only female) | Self-report | 'Yes' response |  |
| Use of hormonal contraceptives (only female) | Self-report | 'Yes' response |  |
| Lactation or breastfeeding (only female) | Self-report | 'Yes' response |  |
| Menstrual cycle (only female) | Reproductive Status Questionnaire for Menstrual Cycle Studies | - |  |
| Color vision deficiency (only for category 2) | Ishihara Test (r | ' Yes ' response |  |
| Chronotype | Ultra-short Munich Chronotype Questionnaire (µMCTQ) | ≤ 2  ≥7 |  |
| Sleep duration | Ultra-short Munich Chronotype Questionnaire (µMCTQ) | < 6  > 9 |  |
| Sleep quality | Pittsburgh Sleep Quality Index, PSQI | >5 |  |
| Smoking | Self-report | >0 |  |
| Substance abuse | Alcohol Use Disorders Identification Test, AUDIT | >7 |  |
| Depressive symptoms | BDI-II | >13 |  |
| High myopia | Self-report from prescription information | < -6 diopters |  |
| High hyperopia | Self-report from prescription information | > +6 diopters |  |
| Transmeridian travel (>2 zones) <1 month prior to first session | Self-report | 'Yes' response |  |
| Shift work <3 months prior to study | Self-report | 'Yes' response |  |
| Current participation in other clinical trials | Self-report | 'Yes' response |  |
| Any ophthalmological or optometric conditions (cataract, glaucoma, retinal detachment, macular conditions, chronic inflammations, eye injuries or operations) | Self-report | Any 'Yes' response |  |
| Any general health concerns or disorders, including heart and cardiovascular, neurological, nephrological, endocrinological and psychiatric conditions | Self-report | Any 'Yes' response |  |
| Any chronic medication affect on sleep | Self-report | Any 'Yes' response |  |
| BMI | Measured height and weight | <18.5  >29.9 | In-person screening session |
| Pregnancy test (only women) | M-Budget pregnancy test | Positive test |  |
| Normal color vision | Cambridge Color Test , Farnsworth Munsell 100 Hue Test | Protan >10, Detran > 10, Tritan > 15, and 100 Hue score > 40 |  |
| Normal best-corrected visual acuity (BCVA) | Landolt C test | Visus < 0.5 |  |
| Ability to understand study language | In-person interaction with the experimenter | Experimenter judgment |  |
| Color vision deficiency | Check by ophthalmologist | Ophthalmologist judgment | Optometry and Physician Screening session |
| Risk of angle-closure glaucoma | Check by ophthalmologist | Ophthalmologist judgment |  |
| Any ophthalmological or optometric conditions (cataract, glaucoma, retinal detachment, macular conditions, chronic inflammations, eye injuries or operations) | Check by ophthalmologist | Ophthalmologist judgment |  |
| Any general health concerns or disorders, including heart and cardiovascular, neurological, nephrological, endocrinological and psychiatric conditions | Check by physician | Physician judgment |  |
| Any chronic medication affect on sleep | Check by physician | Physician judgment |  |
| Drug use (AMP, BZD, COC, MOR/OPI, MTD and THC) | Drug-Screen-Multi 6; nal von Minden | Any positive test | Every study session |
| Alcohol use | Breathalyzer ACE X | >0.05 |  |
| Sleep-wake times in 5 days prior to each experimental session | Actigraphy record and sleep diary | >1 deviation from ±30 minute window sleep and wake-up time |  |
| Ability to follow study instructions | In-person interaction with the experimenter | Experimenter judgment |  |

Table S3) overview light measures of the experimental light conditions

| Condition | Dim light | Bright light |
| --- | --- | --- |
| Luminance [cd/m2] | 13.76 | 294.37 |
| Illuminance [lx] | 8.30 | 100.92 |
| CIE 1931 xy chromaticity [x] | 0.28 | 0.28 |
| CIE 1931 xy chromaticity [y] | 0.29 | 0.29 |
| CIE 1964 x10y10 chromacity: x10 | 0.27 | 0.27 |
| CIE 1964 x10y10 chromacity: y10 | 0.32 | 0.32 |
| CCT (K) - Ohno, 2013 | 9663.259 | 9663.306 |
| CCT (K) - Robertson, 1968 | 9664.5 | 9664.547 |
| S-cone-opic irradiance [mW.m-2] | 4.47 | 74.50 |
| M-cone-opic irradiance [mW.m-2] | 7.53 | 125.55 |
| L-cone-opic irradiance [mW.m-2] | 7.62 | 127.07 |
| Rhodopic irradiance [mW.m-2] | 10.51 | 174.49 |
| Melanopic irradiance [mW.m-2] | 10.59 | 176.49 |
| S-cone-opic EDI [lx] | 6.11 | 101.95 |
| M-cone-opic EDI [lx] | 5.78 | 96.45 |
| L-cone-opic EDI [lx] | 5.23 | 87.25 |
| Rhodopic EDI [lx] | 8.07 | 134.63 |
| Melanopic EDI [lx] | 8.93 | 148.85 |

Table S4) results of the analysis of variance for melatonin AUC, DLMO, and KSS in all participants. P values < 0.05 were considered as significant in bold and p values = 0.06 were presented in italic.

| Outcomes | Predictors | F value | df | p | Effect size (ω_p_^2^) |
| --- | --- | --- | --- | --- | --- |
| Melatonin AUC | Light condition *(Dim and Moderate)* | 3.48 | 1,46 | *0.06* | 0.05 |
|  | Prior light history *(TAT100 lx)* | 0.23 | 1,81 | 0.63 | 0.00 |
| Melatonin DLMO | Prior light history *(TAT100 lx)* | 8.76 | 1, 44 | **0.004** | 0.14 |
| KSS | Light condition *(Dim and Moderate)* | 5.41 | 1,46 | **0.02** | 0.08 |
|  | Prior light history *(TAT100 lx)* | 0.00 | 1,89 | 0.97 | 0.00 |

Table S5) results of the analysis of variance for melatonin AUC, DLMO, and KSS on sex effect. P values < 0.05 were considered as significant in bold and p values = 0.06 were presented in italic.

| Outcomes | Predictors | F value | df | p | Effect size (ω_p_^2^) |
| --- | --- | --- | --- | --- | --- |
| Melatonin AUC | Light condition *(Dim and Moderate)* | 3.41 | 1,45 | *0.06* | 0.05 |
|  | Sex | 0.22 | 1,45 | 0.63 | 0.00 |
|  | Prior light history *(TAT100 lx)* | 0.01 | 1,86 | 0.89 | 0.00 |
|  | Light condition * Sex | 0.43 | 1,52 | 0.51 | 0.00 |
| Melatonin DLMO | Sex | 0.75 | 1,43 | 0.39 | 0.00 |
|  | Prior light history *(TAT100 lx)* | 11.48 | 1,43 | **0.001** | 0.19 |
| KSS | Light condition *(Dim and Moderate)* | 5.42 | 1,45 | **0.02** | 0.09 |
|  | Sex | 0.67 | 1,45 | 0.41 | 0.00 |
|  | Prior light history *(TAT100 lx)* | 0.10 | 1,81 | 0.75 | 0.00 |
|  | Light condition * Sex | 1.14 | 1,53 | 0.28 | 0.00 |

Table S6) results of the analysis of variance for melatonin AUC, DLMO in female participants. P values < 0.05 were considered as significant in bold and p values = 0.06 were presented in italic.

| Outcomes | Predictors | F value | df | p | Effect size (ω_p_^2^) |
| --- | --- | --- | --- | --- | --- |
| Melatonin AUC | Light condition *(Dim and Moderate)* | 3.69 | 1,21 | *0.06* | 0.10 |
|  | Prior light history *(TAT100 lx)* | 0.10 | 1,33 | 0.74 | 0.00 |
|  | Menstrual phase (Follicular and Luteal) | 0.25 | 1,20 | 0.62 | 0.00 |
| Melatonin DLMO | Prior light history *(TAT100 lx)* | 6.92 | 1,21 | **0.01** | 0.20 |
|  | Menstrual phase (Follicular and Luteal) | 8.15 | 1,21 | **0.009** | 0.23 |

Table S7) results of the analysis of variance for melatonin AUC, DLMO, and KSS on season effect. P values < 0.05 were considered as significant in bold.

| Outcomes | Predictors | F value | df | p | Effect size (ω_p_^2^) |
| --- | --- | --- | --- | --- | --- |
| Melatonin AUC | Light condition *(Dim and Moderate)* | 7.99 | 1,45 | **0.006** | 0.13 |
|  | Season | 0.00 | 1,51 | 0.96 | 0.00 |
|  | Prior light history *(TAT100 lx)* | 0.009 | 1,68 | 0.75 | 0.00 |
|  | Light condition * Season | 9.34 | 1,45 | **0.003** | 0.15 |
| Melatonin DLMO | Season | 0.59 | 1,43 | 0.44 | 0.00 |
|  | Prior light history *(TAT100 lx)* | 8.75 | 1,43 | **0.005** | 0.14 |
| KSS | Light condition *(Dim and Moderate)* | 7.03 | 1,44 | **0.01** | 0.11 |
|  | Season | 0.10 | 1,52 | 0.74 | 0.00 |
|  | Prior light history *(TAT100 lx)* | 0.06 | 1,90 | 0.79 | 0.00 |
|  | Light condition * Season | 1.91 | 1,45 | 0.17 | 0.02 |

Table S8) results of the analysis of variance for melatonin AUC, DLMO, and KSS on sex and season effect. P values < 0.05 were considered as significant in bold.

| Outcomes | Predictors | F value | df | p | Effect size (ω_p_^2^) |
| --- | --- | --- | --- | --- | --- |
| Melatonin AUC | Light condition *(Dim and Moderate)* | 8.67 | 1,43 | **0.005** | 0.14 |
|  | Sex | 0.02 | 1,43 | 0.88 | 0.00 |
|  | Season | 0.000.05 | 1,50 | 0.82 | 0.00 |
|  | Prior light history *(TAT100 lx)* | 0.0090.21 | 1,72 | 0.64 | 0.00 |
|  | Light condition * Sex | 0.15 | 1,48 | 0.69 | 0.00 |
|  | Light condition * Season | 10.06 | 1,43 | **0.002** | 0.17 |
|  | Sex * Season | 4.39 | 1,44 | **0.04** | 0.07 |
|  | Light condition * Sex * Season | 3.09 | 1,43 | 0.08 | 0.04 |
| Melatonin DLMO | Sex | 0.73 | 1,41 | 0.39 | 0.00 |
|  | Season | 0.77 | 1,41 | 0.38 | 0.00 |
|  | Prior light history *(TAT100 lx)* | 11.21 | 1,41 | **0.001** | 0.18 |
|  | Sex * Season | 0.24 | 1,41 | 0.62 | 0.00 |
| KSS | Light condition *(Dim and Moderate)* | 7.28 | 1,43 | **0.009** | 0.12 |
|  | Sex | 0.32 | 1,43 | 0.57 | 0.00 |
|  | Season | 0.04 | 1,50 | 0.83 | 0.00 |
|  | Prior light history *(TAT100 lx)* | 0.00 | 1,84 | 0.97 | 0.00 |
|  | Light condition * Sex | 1.49 | 1,49 | 0.22 | 0.00 |
|  | Light condition * Season | 2.05 | 1,43 | 0.15 | 0.02 |
|  | Sex * Season | 0.45 | 1,43 | 0.50 | 0.00 |
|  | Light condition * Sex * Season | 0.78 | 1,43 | 0.37 | 0.00 |

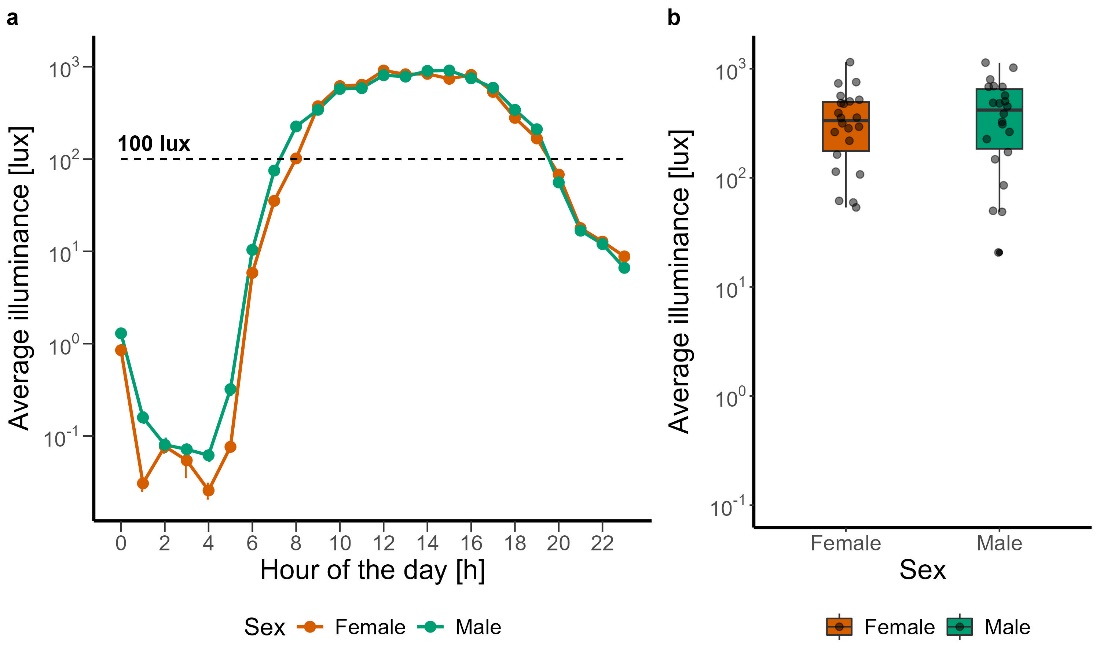

Figure S1. a) average logarithmic illuminance in lux plotted against hour of the day in females (orange) and males (green). b) average logarithmic illuminance in lux in females (orange) and males (green). The box plots display the median (central horizontal line), interquartile range (edges of the box, from first quartile; 25^th^ percentile to third quartile; 75^th^ percentile), the range of minimum and maximum values within 1.5 times the interquartile range from the first and third quartiles (whiskers), and the outliers (solid dots outside of the whiskers). Grey dots represent the individual values of participants.
